# Asymmetric reproductive character displacement and female polymorphism in *Ischnura* damselflies

**DOI:** 10.64898/2026.03.12.711425

**Authors:** Andrea Viviana Ballen-Guapacha, Rosa Ana Sánchez-Guillén

## Abstract

Reproductive Character Displacement (RCD) often occurs when species with mating-related polymorphism come into secondary contact, leading to divergence in reproductive traits. *Ischnura elegans* and *Ischnura graellsii* have formed two independent hybrid zones in Spain where reinforcement has strengthened a mechanical barrier, and RCD has shaped mating-related structures, although reinforcement is asymmetric only in gynochrome females. This study examines the link between asymmetric reinforcement and asymmetric RCD. Using geometric morphometrics, we analyze prothorax shape and size in both female morphs and males, and male caudal appendages, to assess morphological divergence, determine whether gynochrome females show stronger divergence, and test for morphological covariation between male traits involved in the tandem position. Our results reveal consistent patterns of size and shape variation across species and zones: in *I. elegans*, androchromes are larger and resemble males in size, with clear shape differentiation between female morphs that diminishes in hybrid zones. In contrast, *I. graellsii* shows less consistent size differences between males and morphs, and weaker shape differentiation. Our results confirm RCD in prothorax shape in *I. elegans* females from both hybrid zones, but reveal that RCD in prothorax size is asymmetric, occurring only in gynochrome females from the NC hybrid zone. We also detected RCD in the prothorax shape of *I. elegans* males from the NC hybrid zone, extending previous evidence of RCD in male caudal appendages, while morphological covariation between male cerci and the prothorax was limited to size in *I. elegans*. Together, these findings illustrate how hybridization may generate morph-specific patterns of reproductive divergence.

## INTRODUCTION

Reinforcement of reproductive isolation occurs when postzygotic isolation is incomplete, allowing hybrid formation but resulting in sterile or unfit offspring (Coyne & Orr, 1989) or when isolation is complete, but heterospecific matings waste gametes and hinder conspecific fertilization (Howard & Harrison, 1993; Hopkins, 2013). In the first case, selection removes unfit hybrids, while in the second, it favors individuals that avoid costly heterospecific matings. In both scenarios, selection promotes greater phenotypic divergence of reproductive traits in sympatry compared to allopatry, often driving an intensification of trait divergence within one or both species involved (Howard & Harrison, 1993; Cooley, 2007; Pfennig and Pfennig, 2009).

Reproductive Character Displacement (RCD), which refers to the increased phenotypic divergence of reproductive traits in sympatric populations compared to allopatric populations (Schluter, 2000), has been reported in a wide range of taxonomic groups and it is particularly common in species exhibiting mating-related polymorphisms (Pfennig & Murphy, 2002; Rice & Pfennig, 2007). This is because the presence of alternative phenotypes can drive rapid character displacement, as the success of these variants differ when species come into secondary contact (Rice & Pfennig, 2007; Barrett & Schluter, 2008). According to Rice and Pfennig (2007), when species exhibiting mating-related polymorphism are in sympatry with closely related species, selection may favor the morph that is less similar to the competing species [e.g., beetles (Kawano, 2002); tadpoles (Pfennig & Murphy, 2000), butterflies (Hinojosa et al., 2020)]. In plants, RCD has also been invoked to explain patterns of petal color distribution among species, as seen in *Leavenworthia stylosa* (Norton et al., 2015) and *Ruellia* (Tripp et al. 2021).

*Ischnura elegans* and *Ischnura graellsii* are two closely related polymorphic damselfly species. In both, females occur in three female colour morphs: one male-mimic morph (androchrome) and two gynochrome morphs (infuscans and aurantiaca) (Fig. 1) (Cordero 1990; Sánchez-Guillén et al. 2005). This color polymorphism is controlled by a single locus with three alleles showing hierarchical dominance (androchrome>infuscans>aurantiaca) (Cordero-Rivera 1990; Sánchez-Guillén et al. 2005). In *I. elegans*, these morphs consistently exhibit alternative reproductive strategies: androchrome females are less receptive to mating, have lower fecundity, and display more aggressive behavior toward males compared to gynochrome females (Banham 1990; Cordero-Rivera and Sánchez-Guillén 2007; Hammers &Van Gossum 2008; Hammers et al. 2009; Sánchez-Guillén et al. 2017; Cordero-Rivera et al., 2024). In contrast, differences between morphs in *I. graellsii* appear more subtle, with only slightly reduced mating rates in androchrome females (Cordero 1987; 1992). In both species, males typically prefer the gynochrome females, although this preference can shift depending on local morph frequencies (Cordero-Rivera & Andrés-Abad 2001; Cordero-Rivera & Sánchez-Guillén 2007; Sánchez-Guillén et al. 2017).

**Figure 1.**
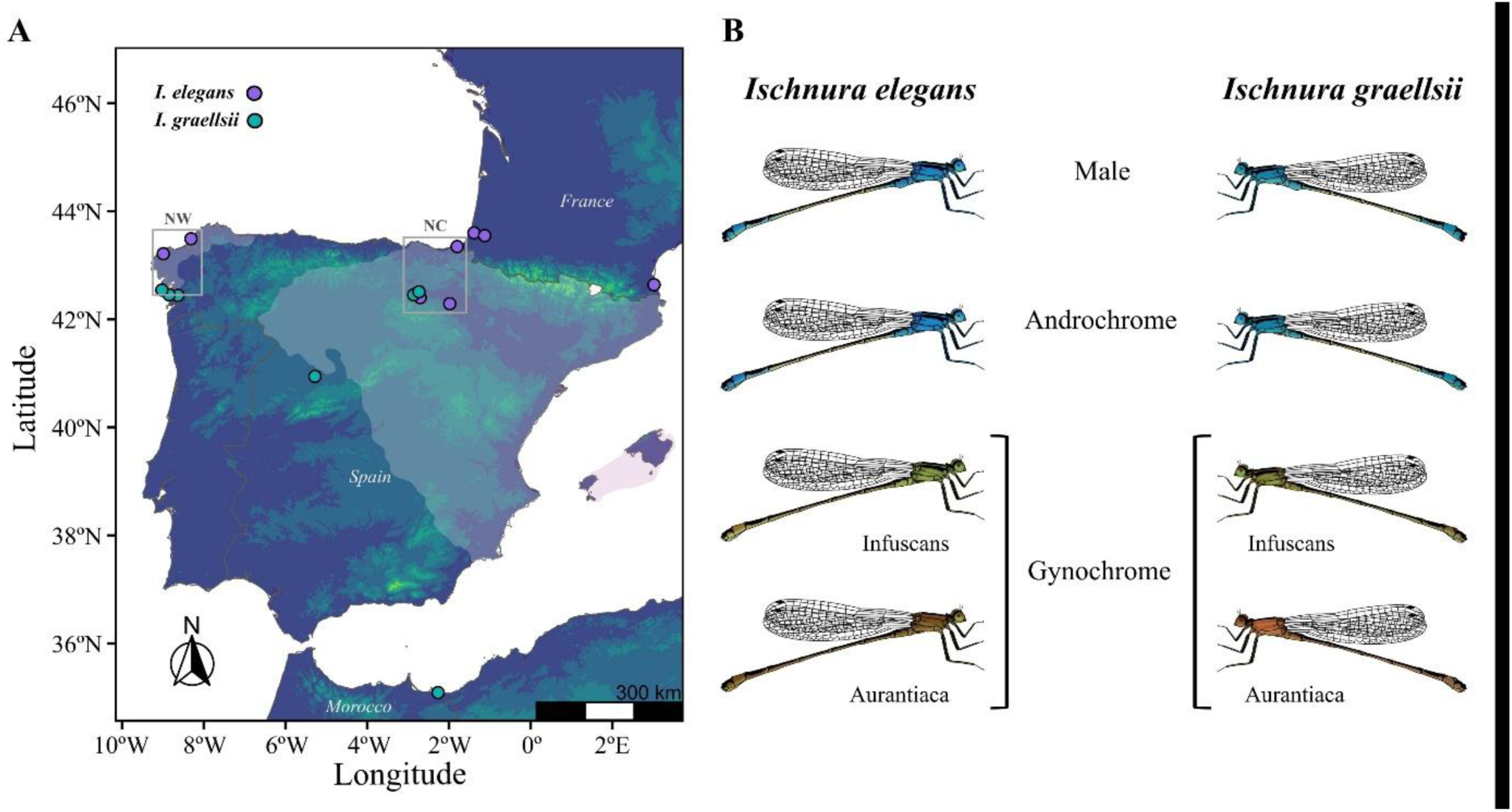
Study sites and color morphs of *Ischnura elegans* and *I. graellsii*. **A)** Map of study localities in allopatry and hybrid zones. Purple dots indicate localities where *I. elegans* was collected, and light blue dots indicate localities where *I. graellsii* was collected. Gray boxes mark the northwest (NW) and the north-central (NC) hybrid zones. Light-yellow regions represent the sympatric distribution of both species. **B**) Illustrations of *I. elegans* and *I. graellsii* showing males and female color morphs. Both species have androchrome females (male-like) and gynochrome females (infuscans and aurantiaca morphs).

These species have evolved in allopatry and have largely maintained separate distributions, with a limited zone of partial sympatry in Spain. More recentrly, however, the range expansion of *I. elegans* has increased their range of overlap, leading to secondary contact and the formation of an extended mosaic hybrid region comprising two unconnected hybrid zones (Fig. 1) (Sánchez-Guillén et al., 2011a; Wellenreuther et al. 2018; Arce-Valdés et al., 2025a). In these hybrid zones, the frequency of female colour morphs differs from that in allopatric regions: the androchrome morph is more common in both species within the hybrid region (Sánchez-Guillén et al., 2005), although high frequencies of the androchrome morph are also reported in allopatric populations in northern Europe (Gosden et al., 2011). In *I. elegans*, morph frequencies are variable at both local and regional scales (Sánchez-Guillén et al., 2005, 2011b; Gosden et al. 2011; Cordero-Rivera et al. 2024), while in *I. graellsii*, they are more stable across its range (Cordero 1990; Andrés et al. 2000; Sánchez-Guillén et al. 2011b). Genetic analyses suggest that in *I. elegans*, divergent selection has acted on the colour locus in the hybrid zone, contrasting with *I. graellsii*, where genetic drift and other selective processes may be involved (Andrés et al. 2000; Sánchez-Guillén et al. 2011b). A similar increase in androchrome frequency has been reported in hybrid zones of North American *Ischnura* species, where it has been linked to male preferences (Johnson 1975).

In these hybrid zones, asymmetric reinforcement of the mechanical barrier has been detected only in crosses between *I. graellsii* males and *I. elegans* females, preventing the initial physical contact during copulation (tandem). This reinforcement has led to RCD in the traits involved in the tandem position, specifically the male cerci of *I. graellsii* and female pronotum of *I. elegans* (Arce-Valdés et al., 2025b; Ballén-Guapacha et al. 2024). Further research by Ordaz-Morales et al. (2025) revealed that the reinforcement of this mechanical barrier is also asymmetric between female morphs of *I. elegans*, (it is present only in the gynochrome morph), while the androchrome morph remains mechanically isolated from heterospecific males both in allopatry and sympatry. This innate isolation in androchrome females has been hypothesized to result from their prothorax morphology, which resembles that of conspecific males and likely prevents effective coupling by heterospecific males. The prothorax plays a key role in the tandem formation, the first step of copulation in odonates, as males grasp females by this structure using their caudal appendages. Differences in prothorax morphology could thus influence mechanical compatibility between individuals (Robertson & Paterson 1982; McPeek et al., 2009; Barnard et al., 2017).

However, it remains unclear whether this asymmetric reinforcement between female morphs is accompanied by morph-specific patterns of RCD in the morphological traits involved in the tandem. In particular, it is unknown whether the stronger reinforcement observed in gynochrome females is reflected in greater morphological divergence of the prothorax relative to androchrome females. In this context, we use geometric morphometric analyses of the reproductive traits involved in the tandem position in both species from the two unconnected hybrid zones to investigate innate differences in female prothorax morphology, the extent of androchrome mimicry of the male prothorax, and whether these innate differences explain the asymmetric reinforcement of the mechanical barrier, which has been detected specifically in gynochrome females when analyzed jointly. Specifically, we predict that (1) female morphs will differ in the shape and/or size of the prothorax, with androchrome females showing greater similarity to conspecific males than gynochrome females, and (2) RCD will be more pronounced in the prothorax of gynochrome morphs pooled together, consistent with the observed asymmetric reinforcement. Additionally, we test whether the shape of the male prothorax and the caudal appendages have coevolved, which would support the idea that mechanical compatibility between sexes operates through a lock-and-key mechanism in these damselflies.

## METHODOLOGY

### Mating system and mechanical isolation during tandem formation

*Ischnura elegans* and *I. graellsii* exhibit a non-territorial mating system where males actively attempt to form tandems with females, but successful mating requires female cooperation (Cordero, 1989; Joop et al., 2006). Copulation starts when the male clasps the female by her prothorax with his caudal appendages, which are considered genital traits because of their functional role in copulation (Brennan, 2016), creating the first contact point, known as the tandem position. After that, the female bends her abdomen, and the male and female genitals come into contact, forming the wheel position.

In sympatric populations, males of both species are equally attracted to conspecific and heterospecific females, indicating a lack of sexual isolation (Sánchez-Guillén et al. 2012). However, mechanical isolation due to incompatibility in forming the tandem position constitutes the main prezygotic reproductive barrier between *I. elegans* and *I. graellsii* (Sánchez-Guillén et al. 2012) as in other damselflies (Sánchez-Guillén et al., 2014; Wellenreuther and Sánchez-Guillén, 2016; Nava-Bolaños et al., 2017; Barnard et al., 2017).

### Geometric morphometric dataset assembly and geometric morphometric analyses

Morphological analyses were based on landmark coordinate data in TPS format from Ballen-Guapacha et al. (2024), which comprised landmark data of the prothorax of 269 females and 329 males of *Ischnura elegans* and *I. graellsii* sampled across allopatry and two independent hybrid zones (NW and NC). To the female dataset, we added new morphometric measurements of the prothorax of 29 additional females in order to balance the representation of androchrome and gynochrome female morphs across the three studied zones (allopatry, NW hybrid zone, and NC hybrid zone). To the male caudal appendages dataset, we added new morphometric measurements of the prothorax of 60 of these males (30 *I. elegans* and 30 *I. graellsii*), that were included in this study, resulting in a combined dataset of the prothorax of 298 females and the caudal appendages and prothorax of 60 males across 15 localities: six allopatric, four from the NW hybrid zone, and five from the NC hybrid zone (Table 1; Fig. 1A). This combined dataset constituted the final dataset used in all morphometric analyses.

**Table 1.**
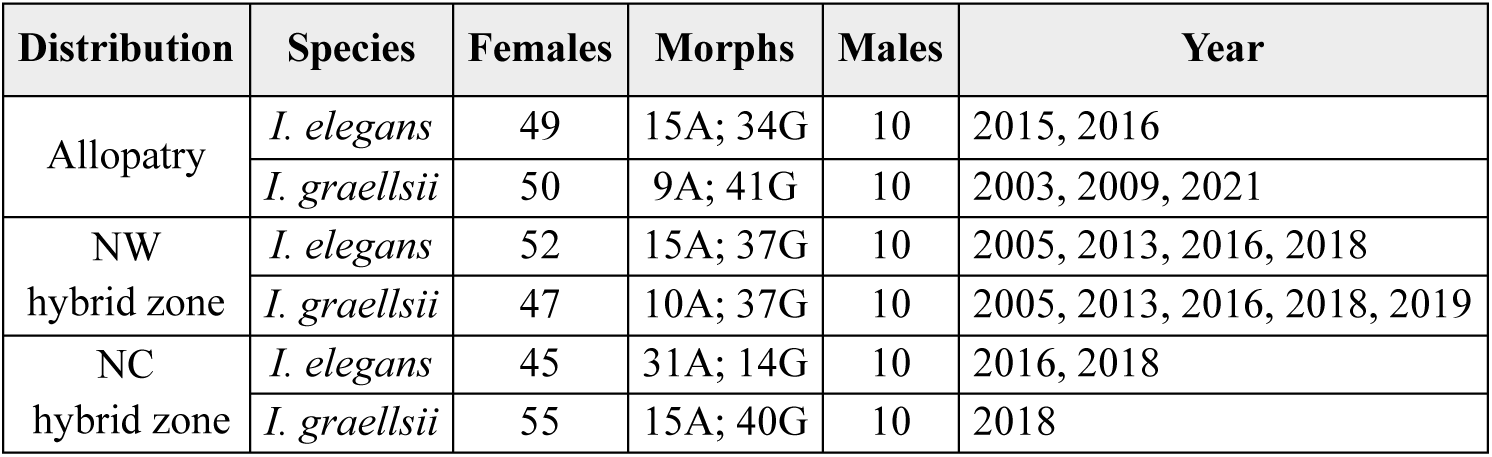
*Ischnura elegans* and *I. graellsii* specimens included in the morphometric analyses. The distribution column indicates the zone where each specimen was collected: allopatry, NW (North-west), and NC (North-central) hybrid zones. N females and N males denote the number of individuals analyzed for the morphometric measurement. A denotes androchrome female morph; G denotes gynochrome female morph.

#### Image acquisition and landmark digitization

Using this final dataset, digital images and landmark coordinated data were obtained for each morphological structure following the procedures described below. Photographs of the posterior and lateral views of male and female prothorax were taken using a Zeiss Stemi 305 stereoscopic microscope with an integrated Axiocam ERc5s camera. The female prothorax was sampled with four landmarks and 21 semi-landmarks on one curve (posterior view) and four landmarks and 19 semi-landmarks on one curve (left lateral view), both surrounding the pronotum. Cartesian coordinates were recorded with tpsDIG2 version 2.04. Data for male caudal appendages were obtained from Ballen-Guapacha et al. (2024). In that study, photographs of the posterior and lateral views of the male caudal appendages were taken using the same microscope and camera setup. The posterior view was sampled using eight landmarks and 80 semi-landmarks distributed along four curves surrounding each cercus. The lateral view was sampled with seven landmarks and 36 semi-landmarks placed along two curves covering the cercus and paraproct. Cartesian coordinates were also recorded using tpsDIG2 version 2.04, and geometric morphometric analyses, including a Generalized Procrustes Analysis (GPA) (Rohlf and Slice, 1990) and thin-plate spline analysis, were conducted using the Geomorph package version 4.0.4 (Adams et al., 2022).

In this study, a GPA (Rohlf and Slice, 1990) was performed to align all landmark configurations by removing variations due to translation, orientation, and scale, followed by thin-plate spline analysis (Bookstein, 1991) using the Geomorph package 4.0.4 (Adams et al., 2022). Semilandmarks were aligned using the Bending Energy minimization criteria (Gunz and Mitteroecker, 2013). Centroid size (CS) was calculated for each configuration as a size estimator (Dryden & Mardia 2016; Klingenberg, 2020). To statistically assess shape differences among males, androchrome females, and gynochrome females, we performed a Procrustes ANOVA including CS and its interaction with Morph (male, androchrome and gynochrome) as predictors, while differences in CS were evaluated using linear models as a function of Morph. Shape variation was analyzed using Principal Components Analysis (PCA) to describe morphological changes, also performed with Geomorph version 4.0.4 (Adams et al., 2022).

### Female morph differentiation and androchrome mimicry of male prothorax morphology

To determine whether androchrome and gynochrome female morphs vary in the shape and/or CS of the prothorax, a comprehensive morphometric analysis was conducted. Additionally, we assessed whether the shape and/or the CS of the prothorax of the androchrome female morph more closely resembles that of males compared to the gynochrome female morph.

To this end, we performed the statistical analyses in the following manner. Firstly, to assess the statistical hypotheses investigating differences among males, androchrome and gynochrome females in shape variation for the set of (Procrustes-aligned) coordinates, we performed a Procrustes ANOVA model using the ‘procD.lm’ function from the Geomorph package version 4.0.4 (Adams et al., 2022). This function calculates the Procrustes (morphological) distance variance explained by each factor in the model, which were the CS, geographic group (allopatric zone, NW hybrid zone, or NC hybrid zone) and their interaction. When the geographic group factor was statistically significant, we did a pairwise evaluation using the ‘permudist’ function from the ‘Morpho’ package version 2.8 (Schlager, 2017). This function compares the distance between the means of two groups with the distances obtained when observations are randomly assigned to groups (here 1,000 permutations) (Schlager, 2017).

In a similar manner, to assess the statistical hypotheses investigating differences among males, androchrome and gynochrome female morphs in CS variation we used the function ‘lm.rrpp’ from the RRPP package (Collyer & Adams, 2018, 2021) to perform linear models. When the geographic group factor showed statistical significance, we did pairwise comparisons to detect which groups were statistically different using the ‘pairwise’ function. Differences were estimated as Euclidean distance between least-squares means to each group and the significance of distances was assigned using the permutation procedure previously described.

### Inference of reproductive character displacement

We assessed whether the asymmetric RCD in the prothorax morphology between species previously reported by Ballén-Guapacha et al. (2024) is stronger in one female morph, morph-specific, or also detectable in males. Thus, we measured RCD of the shape and CS of the prothorax in both female morphs and males, using the key criterion that the interspecific morphological distance between these reproductive characters is significantly greater in sympatry than in allopatry (Slatkin, 1980; Liou & Price, 1994). Additionally, when this pattern was detected, we performed intraspecific comparisons to determine whether trait values differed significantly between sympatric and allopatric populations in one or both species. To this end, we first calculated the interspecific morphological distance between androchrome *I. elegans* and androchrome *I. graellsii* in the allopatric zone and compared this distance with distances from hybrid zones. A significantly larger morphological distance in the hybrid zone, compared to the allopatric zone, indicated evidence of RCD. We repeated this process for gynochrome female morph and for males. Next, in cases of RCD, we compared intraspecific morphological distances to determine if one or both species had significantly different sizes in allopatry versus sympatry. RCD was considered unilateral if the significant change in morphology occurred in only one species or if both species changed in the same direction but with a larger magnitude in one species. RCD was considered bilateral if both species showed significant changes in morphology, with each species changing in opposite directions (one larger, the other smaller). Lastly, we compared the occurrence and relative magnitude of RCD among androchrome female morph, gynochrome female morph, and males to understand the extent and variation of this phenomenon across different morphs and sexes. To this end, we performed the statistical analyses in the following manner. We used a Procrustes ANOVA model via the ‘procD.lm’ function in the Geomorph package version 4.0.4 (Adams et al., 2022) to evaluate the statistical hypotheses regarding RCD patterns of shape variation for the set of Procrustes-aligned coordinates. When the geographic group factor was found to be statistically significant, we used the R base package (R core team, 2023) to perform contrast tests evaluating whether interspecific morphological distances in each hybrid zone differed significantly from the allopatric reference. When distances were significantly larger in sympatry with respect to allopatry, we supplemented this analysis with an intraspecific pairwise evaluation using the ‘permudist’ function from the Morpho package version 2.8 (Schlager, 2017). Similarly, to evaluate the statistical hypotheses regarding RCD patterns in CS, we used the ‘lm.rrpp’ function from the RRPP package (Collyer & Adams, 2018, 2021) to perform linear models, and when the group factor was statistically significant, we used the R base package (R core team, 2023) to perform contrast tests evaluating whether interspecific distances in each hybrid zone differed significantly from the allopatric reference. When distances were significantly larger in sympatry with respect to allopatry, we conducted intraspecific pairwise comparisons using the ‘pairwise’ function to determine which groups were significantly different (see Ballén-Guapacha et al. 2024 for a full description of the statistical framework).

### Morphological integration between male caudal appendages and prothorax

We examined correlations between the shape and CS of male caudal appendages and the male prothorax in both species. If these structures are developmentally or functionally integrated, the magnitude of change in caudal appendages should correlate with changes in the prothorax.

To test this, we assessed correlations in both shape and CS separately for *I. elegans* and *I. graellsii* analyzing individuals from allopatric populations as a reference for non-hybrid conditions, as well as from sympatric populations within both hybrid zones (NW and NC) to evaluate whether patterns of morphological integration are maintained or altered under conditions of hybridization and reinforcement. We conducted a two-block partial least squares analysis (PLS) on Procrustes shape variables and CS variables using the ’two.b.pls’ function from the Geomorph package version 4.0.4 (Adams et al., 2022). All analyses were conducted with the statistical software R (version 4.3.0, R Core Team, 2023).

## RESULTS

### Female morph differentiation and androchrome mimicry of male prothorax morphology

The results from the posterior view were largely consistent with those from the lateral view. To avoid redundancy, we present the results from the lateral view in the main text, and provide the corresponding posterior view results in the Supplementary Material.

#### Prothorax shape variation

In *I. elegans*, to assess differences in the prothorax shape (lateral view) between males and female morphs in the three zones (allopatry, NW and NC hybrid zones), we performed procrustes ANOVAs. These analyses showed statistically significant differences between males and female morphs in allopatry (F(2, 56) = 3.681, P < 0.001), in the NW (F(2, 59) = 1.940, P = 0.046) and in the NC (F(2, 52) = 2.518, P = 0.014) hybrid zones (Fig. 2A–C; Supplementary Table 1). Post hoc pairwise tests showed statistically significant differences between males and the gynochrome females in (D = 0.21, P < 0.001), as well as in the NW (D = 0.09, P < 0.005) and in the NC (D = 0.13, P < 0.001) hybrid zones (Supplementary Table 2). In contrast, no significant differences were found between males and androchrome females in any of the zones (allopatry: D = 0.07, P = 0.393; NW: D = 0.07, P = 0.067; NC: D = 0.04, P = 0.409; Supplementary Table 2), indicating a close similarity in prothorax shape between males and androchrome females across all geographic contexts. Pairwise tests between androchrome and gynochrome female morphs showed statistically significant differences in allopatry (D = 0.15, *P* < 0.001) and in the NC hybrid zone (D = 0.09, P < 0.001), but not in NW hybrid zone (D = 0.04, P = 0.467), suggesting reduced differentiation between female morphs in sympatry in the NW hybrid zone.

**Figure 2.**
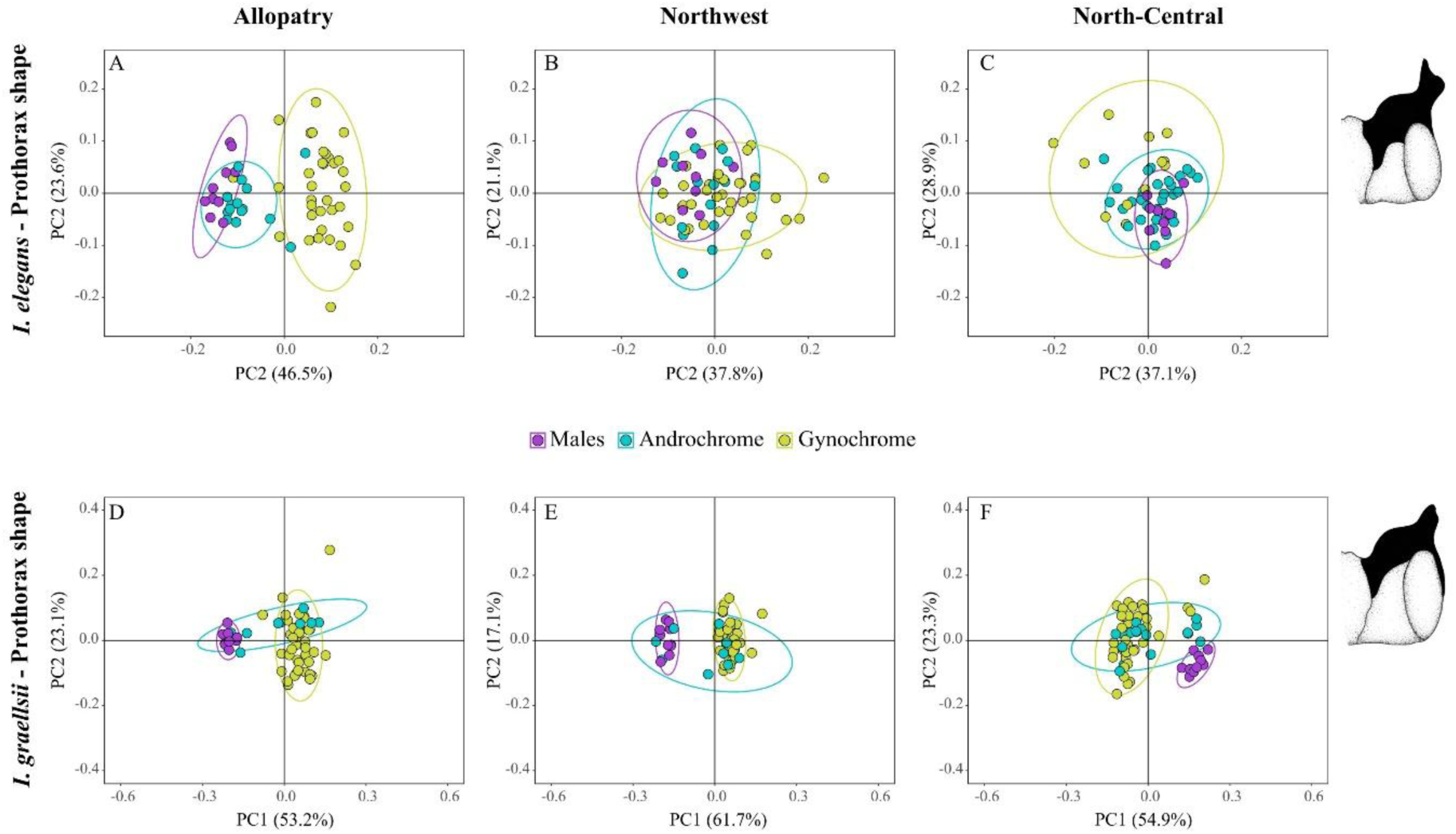
Principal Component Analysis (PCA) of prothorax shape in *Ischnura elegans* and *I. graellsii* (lateral view). Panels **A–C** show *I. elegans*, and panels **D–F** show *I. graellsii* across different geographic zones (allopatry, NW, and NC hybrid zones). Each panel represents morphological variation captured by the first two principal components (PC1 and PC2). Percentages of variance explained by each component are indicated for each species and zone: 70.1%, 58.9%, and 66.0% for *I. elegans* in allopatry, NW, and NC zones, respectively; and 76.3%, 78.8%, and 78.2% for *I. graellsii* in allopatry, NW, and NC zones, respectively.

In *I. graellsii*, procrustes ANOVA to evaluate differences in the shape (for lateral view) between males and female morphs revealed statistically significant differences in allopatry and both hybrid zones (Allopatry: F(2, 57) = 23.809, P < 0.001; NW: F(2, 54) = 20.616, P < 0.001; and NC: F(2, 62) = 13.660, P < 0.001; Fig. 2D–F; Supplementary Table 1). Post hoc pairwise tests showed statistically significant differences between males and gynochrome females in the three zones (Allopatry: D = 0.24, P < 0.001; NW: D = 0.21, P < 0.001; NC: D = 0.23, P < 0.001; Supplementary Table 2). Similarly, significant differences between males and androchrome females were found in all zones (allopatry: D = 0.21, P < 0.001; NW: D = 0.18, P = 0.002; NC: D = 0.21, P < 0.001). In contrast, comparisons between the androchrome and gynochrome females, showed statistically significant differences only in allopatry (D=0.10, P =0.037; Supplementary Table 2), but not in the NW (D = 0.06, P = 0.131) or NC (D = 0.03, P = 0.898) hybrid zones.

Results from the posterior view are provided in Supplementary Tables 3–4, and Supplementary Fig. 2.

#### Prothorax size variation

In *I. elegans* ANOVA analyses revealed statistically significant differences in CS between males and female morphs in allopatry (F(2, 56) = 36.242, P < 0.001, NW (F(2, 59) = 23.369, P < 0.001, and NC hybrid zones (F(2, 52) = 21.261, P < 0.001; Fig. 3; Supplementary Table 5). Post hoc analyses showed that males were not significantly larger than androchrome females in allopatry (Z = 0.782, P = 0.241), but were significantly larger in the NW (Z = 2.806, P = 0.002) and NC hybrid zones (Z = 2.667, P = 0.002). In contrast males were significantly larger than gynochrome females in all three zones (allopatry: Z = 3.776, P = 0.001; NW: Z = 3.484, P = 0.001; NC hybrid zone: Z = 3.691, P = 0.001; Fig. 3; Supplementary Table 6). Between female morphs, androchrome females were significantly larger than gynochrome females in all three zones (allopatry: Z = 3.176, P < 0.001; NW: Z = 2.896; NC: Z = 2.368, P = 0.002) (Supplementary Table 6).

**Figure 3.**
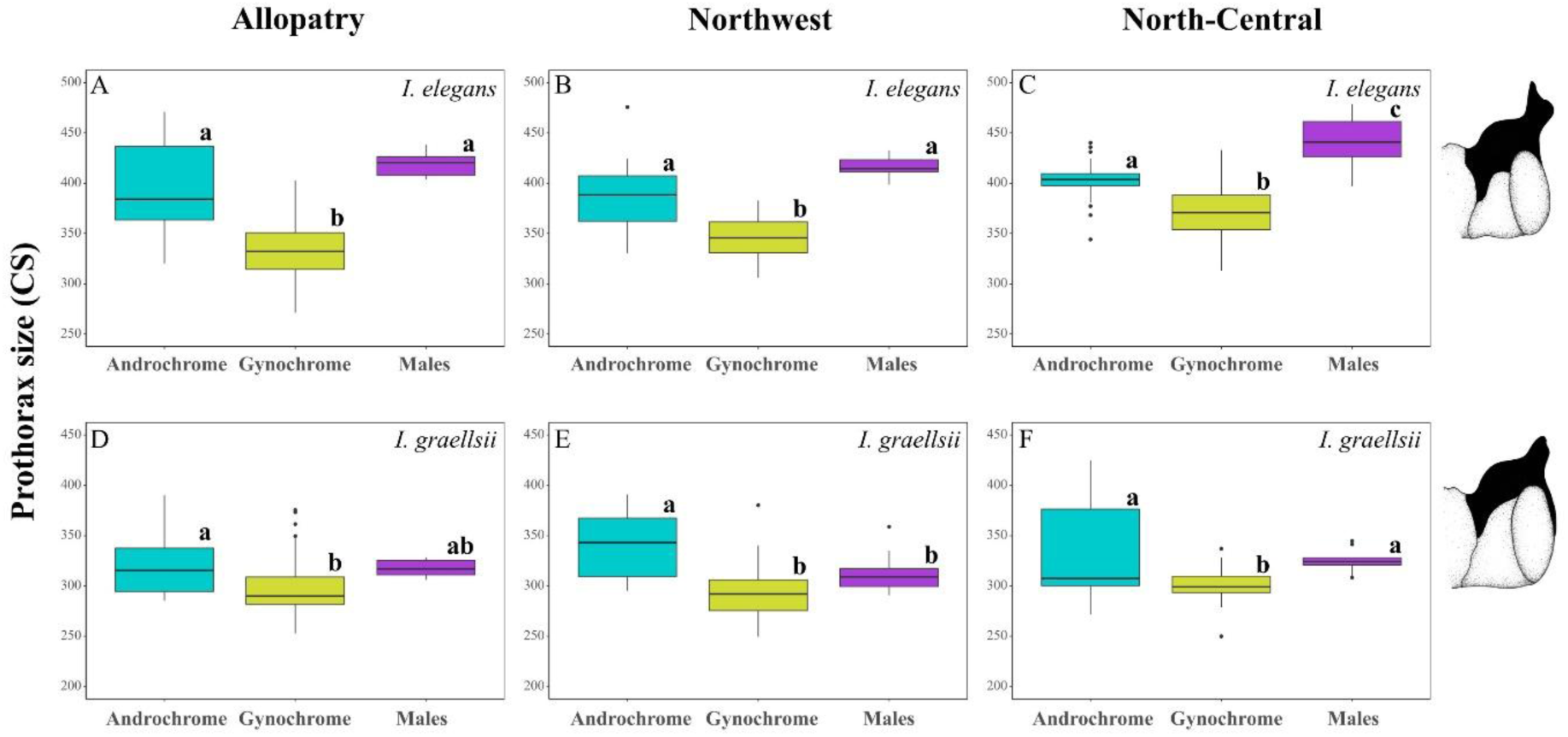
Boxplots of prothorax CS (lateral view) in males and female morphs of *Ischnura elegans* and *I. graellsii*. Panels **A–C** show *I. elegans*, and panels **D–F** show *I. graellsii*. For each species, comparisons are made across geographic zones (allopatry, NW, and NC hybrid zones). Within each plot, males and both female morphs (androchrome and gynochrome) are displayed. Boxes represent interquartile ranges (25th–75th percentiles), horizontal lines indicate medians, and whiskers extend to minimum and maximum values.

In *I. graellsii* ANOVA revealed statistically significant differences in CS in allopatry (F(2, 57) = 3.539, P = 0.041) and both hybrid zones (NW: F(2, 54) = 11.676, P < 0.001; NC: F(2, 62) = 10.078, P = 0.002) (Fig. 3; Supplementary Table 5). Post hoc analyses showed that males were significantly larger than the androchrome females in the NW zone (Z = 1.687, P = 0.041) but not in the allopatry (Z = -0.987, P = 0.820), nor in the NC hybrid zone (Z = 0.116, P = 0.477) (Supplementary Table 6). Males were significantly larger than gynochrome females only in the NC hybrid zone (Z = 1.833, P = 0.026) but not in allopatry (Z = 1.514, P = 0.060) nor in NW hybrid zone (Z = 1.164, P = 0.128). Between female morphs, androchrome females were significantly larger than gynochrome females in all zones (allopatry: Z = 1.686, P = 0.045; NW: Z = 3.272, P < 0.001; NC: Z = 2.933, P < 0.001; Supplementary Table 6).

Results from the posterior are provided in Supplementary Tables 7-8, and Supplementary Fig. 3.

### Inference of reproductive character displacement

The results from the posterior view were largely consistent with those from the lateral view. To avoid redundancy, we present the lateral view results in the main text and provide the posterior view results in Supplementary Tables 10, 11 and 14, and Supplementary Figs. 5 and 6.

#### Shape variation of prothorax

In males, PCA plots (lateral view) showed a clear separation between species in the three zones (see Supplementary Fig. 4A-C). Procrustes ANOVA showed statistically significant differences between *I. elegans* and *I. graellsii* in both hybrid zones (NW: F(1, 18) = 9.592, P < 0.001; NC: F(1, 18) = 9.482, P < 0.001; Supplementary Table 9), but not in allopatry (F(1, 18) = 1.801, P = 0.117). Contrast tests indicated that interspecific distances in both hybrid zones (NW: D = 0.318; NC: D = 0.318; Fig. 4A) were significantly larger than the interspecific distance in allopatry (D = 0.317; Fig. 4A) (NW: t = 2.249, d.f. = 27, P = 0.033; NC: t = 2.271, d.f. = 27, P = 0.031; Supplementary Tables 10 and 11). Therefore, the variance in distance supported the RCD in both hybrid zones (see Table 2). We conducted intraspecific comparisons in both species to investigate whether the RCD was unilateral or bilateral (i.e., due to changes in one or both species). Procrustes ANOVA showed statistically significant differences among males of *I. elegans* from the three zones (F = 4.144, P < 0.001; Fig. 4C; Supplementary Table 12). Post hoc pairwise tests showed statistically significant differences between males from the allopatric zone and the NC hybrid zone (NW: P < 0.141; NC: P < 0.001); Fig. 4C; Supplementary Table 13). While in *I. graellsii* Procrustes ANOVA showed no significant differences in the in males from the three zones (F = 1.421, P = 0.257; Fig. 4C; Supplementary Table 12). Therefore, we detected RCD for the males of *I. elegans* in the NC hybrid zone, while for *I. graellsii* we did not detect RCD in any of the two hybrid zones (see Table 2).

**Figure 4.**
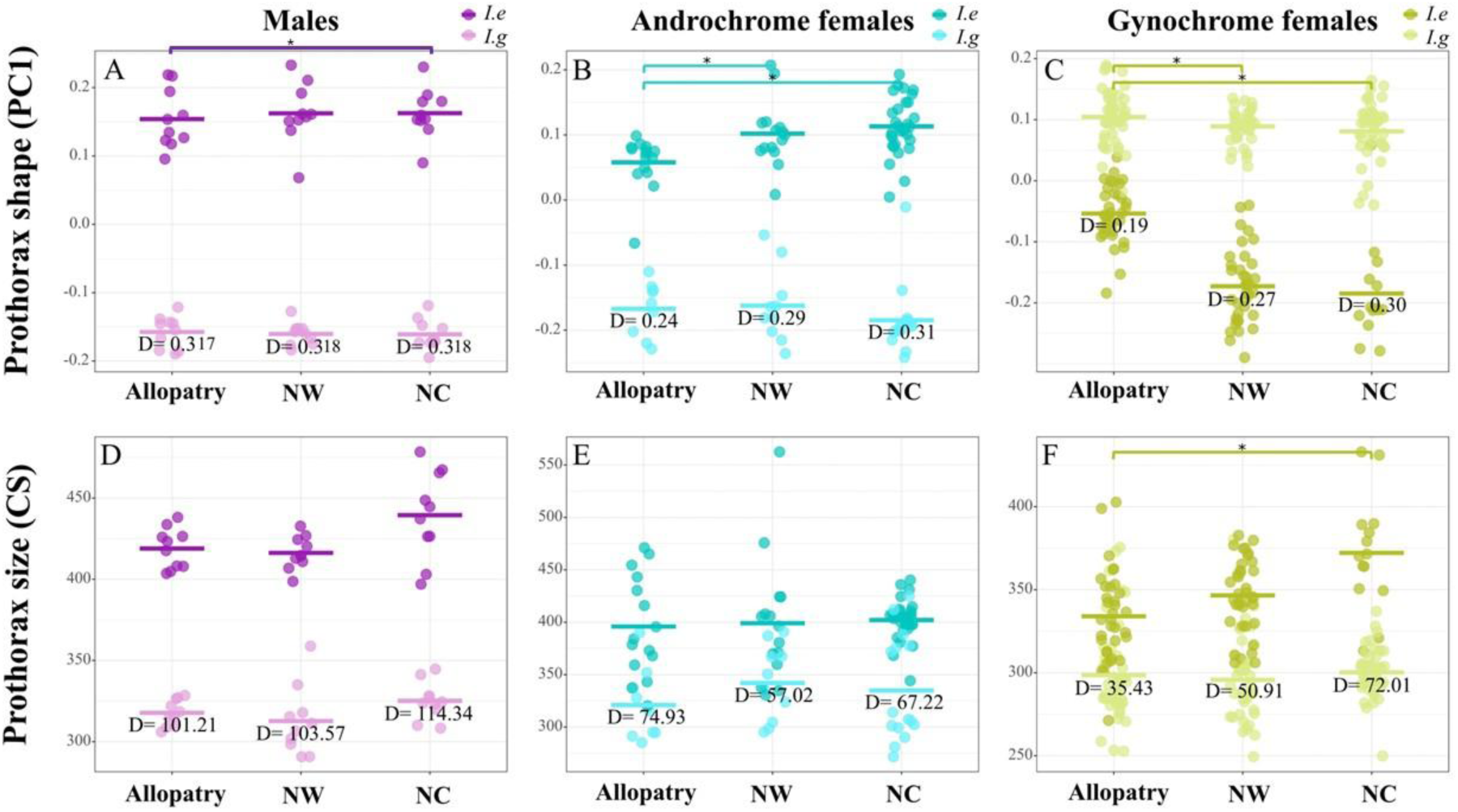
Evidence of reproductive character displacement (RCD) in prothorax shape and CS (lateral view) of *Ischnura elegans* and *I. graellsii*. Panels **A, D** show males, panels **B, E** show androchrome female morphs, and panels **C, F** show gynochrome female morphs across geographic zones (allopatry, NW, and NC hybrid zones). Median values of prothorax shape and CS are displayed. Asterisks above plots indicate statistically significant differences (**P < 0.05**).

**Table 2.**
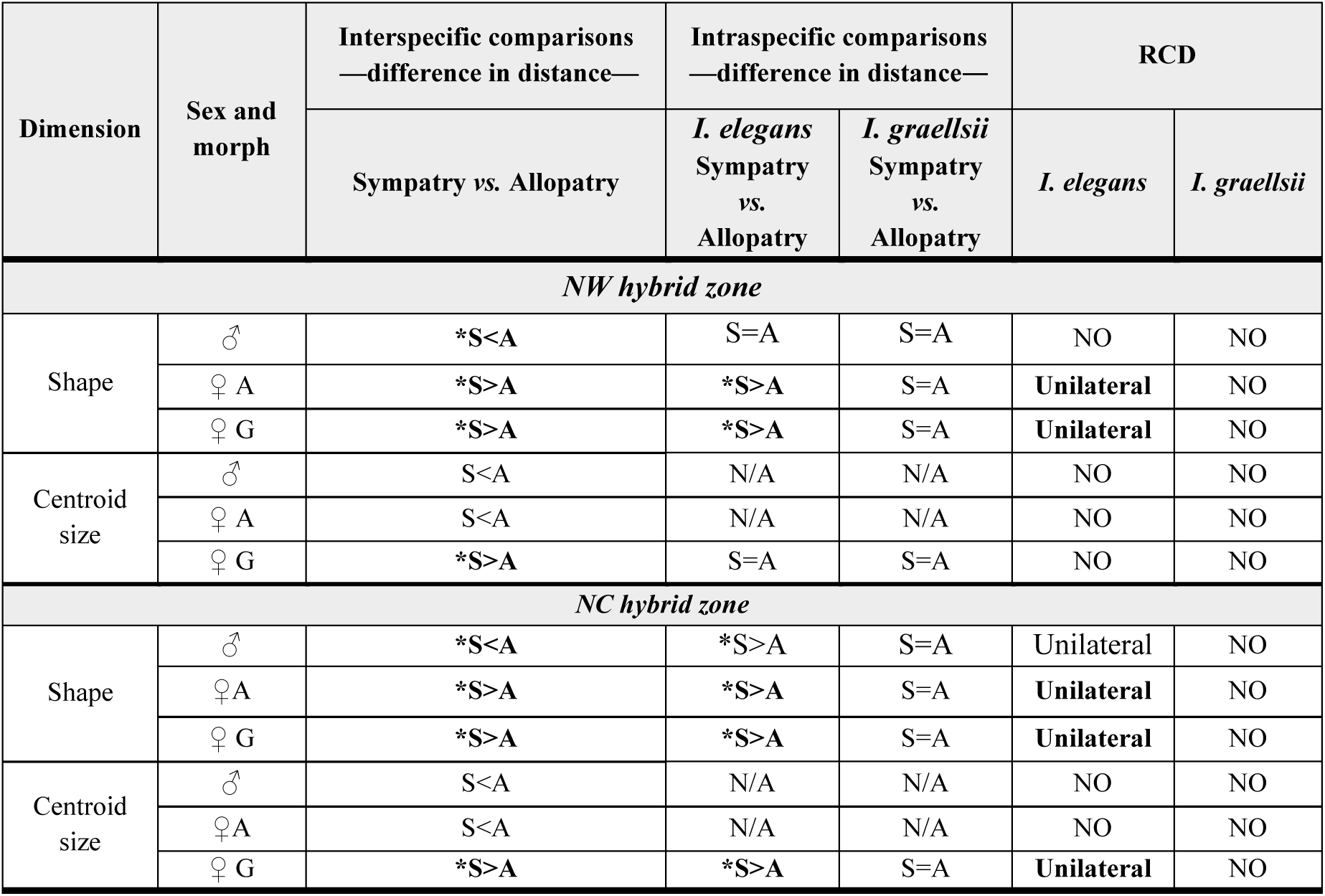
RCD based on the lateral view of the prothorax in *I. elegans* and *I. graellsii* from both hybrid zones. The first two columns indicate the view analyzed (shape or CS), sex (male or females), and female morph (androchrome or gynochrome). The third column shows interspecific comparisons: a significantly larger morphological distance in the hybrid zone than in allopatry (interspecific distance in sympatry – interspecific distance in allopatry > 0) provides support for RCD. The fourth column presents intraspecific comparisons, performed when RCD was supported, to determine RCD, and whether displacement (intraspecific distance in sympatry>intraspecific distance in allopatry) occurred or not, in one or both species. The final column classifies RCD as unilateral (significant change in only one species, or both change in the same direction but with a larger magnitude in one) or bilateral (both species show significant change in opposite directions: one increases, the other decreases). Asterisks indicate statistically significant comparisons. N/A = not applicable.

In androchrome females, PCA plots (lateral view) showed a clear separation between species in the three zones (Supplementary Fig. 4D-F). Procrustes ANOVA showed statistically significant differences between *I. elegans* and *I. graellsii* in allopatry (F (1, 22) = 13.000, P < 0.001), and in both hybrid zones (NW: F(1, 23) = 21.752, P < 0.001; NC: F(1, 44) = 39.272, *P* < 0.001; Supplementary Table 9). Contrast tests indicated that interspecific distances in both hybrid zones (NW: D = 0.29; NC: D = 0.31; Fig. 4B) were significantly larger than the distance in allopatry (D = 0.24; Fig. 4B) (NW: t = 4.674, d.f. = 42, P < 0.001; NC: t = 5.200, d.f. = 42, *P* < 0.001; Supplementary Tables 10 and 11). Thus, distance variance supports the RCD in both hybrid zones (see Table 2). We conducted intraspecific comparisons in both species to investigate RCD and whether it was unilateral or bilateral. Procrustes ANOVA showed statistically significant differences among androchrome female morphs of *I. elegans* from the three zones (F = 5.058, *P* < 0.001; Fig. 4C; Supplementary Table 12). Post hoc pairwise tests showed statistically significant differences between the androchrome female morph from the allopatric zone and both hybrid zones (NW: *P* < 0.001; NC: *P* < 0.001); Fig. 4C; Supplementary Table 13). While in *I. graellsii* Procrustes ANOVA showed no significant differences in the gynochrome female morphs from the three zones (F = 0.491, P = 0.834; Fig. 4C; Supplementary Table 12). Therefore, we detected RCD for the androchrome female morph of *I. elegans* in both hybrid zones, while for *I. graellsii* we did not detect RCD in any of the two hybrid zones (see Table 2).

In gynochrome females, PCA plots (lateral view) showed a clear separation between species in the three zones (Supplementary Fig. 4G-I). Procrustes ANOVA showed statistically significant differences between *I. elegans* and *I. graellsii* in allopatry (F(1, 73) = 28.418, *P* < 0.001), and in both hybrid zones (NW: F(1, 72) = 44.429, P < 0.001; NC: F(1, 52) = 9.218, P < 0.001; Supplementary Table 9). Contrast tests indicated that interspecific distances in both hybrid zones (NW: D = 0.27; NC: D = 0.30; Fig. 4C) were significantly larger than the distance in allopatry (D= 0.19; Fig. 4C). (NW: t = 9.432, d.f. = 108, P < 0.001; NC: t = 8.506, d.f. = 108, P < 0.001; Supplementary Tables 10 and 11). Thus, interspecific morphological distances were smaller in the allopatric zone than in either hybrid zones, supporting RCD in both hybrid zones. We conducted intraspecific comparisons in both species to investigate he RCD and whether it was unilateral or bilateral. Procrustes ANOVA showed statistically significant differences between gynochrome female morph of *I. elegans* from the three zones (F = 11.466, P < 0.001; Fig. 4C; Supplementary Table 12). Post hoc pairwise tests showed statistically significant differences between the gynochrome female morphs from the allopatric zone and both hybrid zones (NW: P < 0.001; NC: P < 0.001); Fig. 4C; Supplementary Table 13). In *I. graellsii*, procrustes ANOVA showed statistically significant differences among gynochrome female morphs of *I. graellsii* from the three zones (F = 2.601, P = 0.004; Fig. 4C; Supplementary Table 12). Post hoc pairwise test showed non-statistically significant differences between the gynochrome female morphs from the allopatric zone and the NW hybrid zone (P = 0.219; Supplementary Table 13) and between gynochrome female morphs from allopatry and the NC hybrid zone (P = 0.072). Therefore, we detected RCD for the gynochrome female morph of *I. elegans* in both hybrid zones (NW and NC), while for *I. graellsii* we did not detect RCD in any of the two hybrid zones (see Table 2).

#### Size variation of prothorax

In males, ANOVA showed statistically significant differences between *I. elegans* and *I. graellsii* in allopatry (F = 464.9, P < 0.001), and in both hybrid zones (NW: F = 197.41, P < 0.001; NC: F = 151.42, P < 0.001; Supplementary Table 15). However, contrast tests indicated that interspecific distances in both hybrid zones (NW: D = 103.57; NC: D = 114.34; Fig. 4D; Supplementary Table 16) were not significantly larger than the distance in allopatry (D = 101.21; Fig. 4D; Supplementary Table 16). (NW: t = 0.563, d.f. = 27, P = 0.578; NC: t = 1.000, d.f. = 27, P = 0.326; Supplementary Table 11). Thus, distance variance did not support the RCD in either hybrid zone (see Table 2).

In androchrome females, ANOVA showed statistically significant differences between *I. elegans* and *I. graellsii* in allopatry (F = 16.425, P < 0.001), and in both hybrid zones (NW: F = 7.264, P = 0.014; NC: F = 43.043, P < 0.001; Supplementary Table 15). However, contrast tests indicated that interspecific distances in both hybrid zones (NW: D = 57.02; NC: D = 67.22; Fig. 4E; Supplementary Table 16) were not significantly larger than the distance in allopatry (D = 74.93; Fig. 4E; Supplementary Table 16) (NW: t = 0.640, d.f. = 42, P = 0.526; NC: t = 0.134, d.f. = 42, P = 0.894; Supplementary Table 11). Therefore, the variance in distance did not support the RCD for the androchrome female morph in any of the hybrid zones (see Table 2).

In gynochrome females, ANOVA showed statistically significant differences between *I. elegans* and *I. graellsii* in allopatry (F = 28.440, P < 0.001), and in both hybrid zones (NW: F = 82.382, P < 0.001; NC: F = 114.670, P < 0.001 Supplementary Table 15). Contrast tests indicated that interspecific distances in both hybrid zones (NW: D = 50.91; NC: D = 72.01; Fig. 4F; Supplementary Table 16) were significantly larger than the distance in allopatry (D = 35.43; Fig. 4F; Supplementary Table 16) (NW: t = 2.114, d.f. = 108, P = 0.037; NC: t = 4.779, d.f. = 108, P < 0.001; Supplementary Table 11). Thus, distance variance supported the RCD in both hybrid zones. We conducted intraspecific comparisons to investigate RCD and whether it was unilateral or bilateral. ANOVA showed statistically significant differences among gynochrome females of *I. elegans* from the three zones (F = 10.384, P < 0.001; Supplementary Table 17). Post hoc pairwise tests showed statistically significant differences between gynochrome females of *I. elegans* only from allopatry and the NC hybrid zone (P < 0.001); Fig. 4F; Supplementary Table 18). However, in the gynochrome females of *I. graellsii*, ANOVA showed non statistically significant differences among zones (F(= 0.331; P = 0.727; Supplementary Table 17). Therefore, we detected RCD in the gynochrome females of *I. elegans* from the NC hybrid zone but not in *I. graellsii* from any hybrid zone (see Table 2). Results of the posterior view are shown in the Supplementary Tables 16 and 19 and the Supplementary Fig. 6.

### Correlation between male caudal appendages and male prothorax

We investigated whether there is a correlation (in terms of shape and CS) between male caudal appendages and male prothorax in allopatric populations, as well as in both hybrid zones. We did not detect a significant correlation between the shape (lateral and posterior views) of the caudal appendages and the prothorax of the *I. elegans* males (lateral view: r = 0.522; P = 0.354; posterior view: r = 0.619; P = 0.281; Supplementary Fig. 7A-B; Supplementary Table 20) or *I. graellsii* males (lateral view: r = 0.529; P = 0.434; posterior view: r = 0.482; P = 0.291) (Supplementary Fig. 7C-D; Supplementary Table 20). However, we detected a significant (although marginal for the lateral view) positive correlation between the CS (lateral and posterior views) of the caudal appendages and the prothorax of the *I. elegans* males (lateral view: r = 0.339; P = 0.077; posterior view: r = 0.429; P = 0.013; Fig. 5A-B; Supplementary Table 20) while this correlation was not detected for any views in *I. graellsii* males (lateral view: r = -0.137; P = 0.474; posterior view: r = 0.256; P = 0.175) (Fig. 5C-D; Supplementary Table 20).

**Figure 5.**
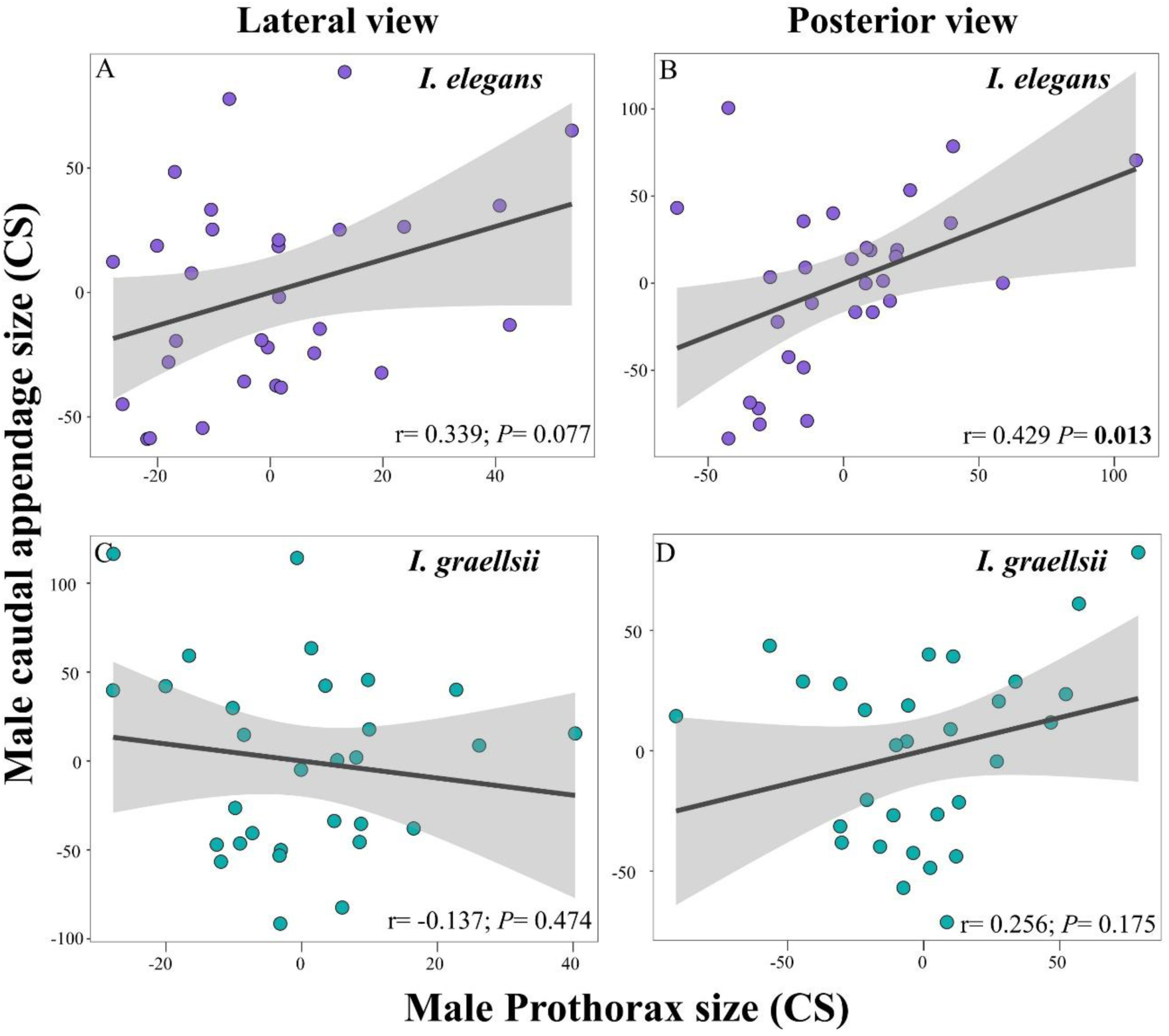
Correlation between cercus and prothorax CS (lateral and posterior views) in males of *Ischnura elegans* and *I. graellsii*. Lines represent linear regression fits, with shaded areas indicating 95% confidence intervals. Correlation coefficients (r) and corresponding significance levels are shown in each plot.

## DISCUSSION

Our study provides new evidence regarding the selective pressures shaping female morphs of *I. elegans* and *I. graellsii* in allopatry and within hybrid zones.

### Morphological basis of male mimicry in androchrome females

In *I. elegans,* gynochrome females differ from males in both prothorax shape and size, and notably, consistently differ from androchrome females, a pattern less evident in *I. graellsii*. These findings suggest that in *I. elegans*, gynochrome females experience distinct selective pressures, potentially linked to their higher exposure to male mating attempts, as males typically prefer this morph (Cordero-Rivera & Andrés-Abad 2001; Cordero-Rivera & Sánchez-Guillén 2007; Sánchez-Guillén et al. 2013, 2017). Importantly, our results show that androchrome females of *I. elegans* exhibit a prothorax shape that closely resembles that of males in both allopatry and sympatry, and a prothorax size that also resembles males, particularly in allopatry and the NC hybrid zone. This morphological resemblance is consistent with the male-mimicry hypotheses (Robertson 1985; Hinnekint 1987; Sherratt 2001) which propose that mimicry may reduce male harassment. As such, androchrome females can adopt an alternative reproductive strategy by being less prone to mate compared to gynochrome females (Cordero-Rivera & Sánchez-Guillén 2007; Hammers & Van Gossum 2008; Hammers et al. 2009; Cordero-Rivera et al., 2024). Additionally, androchrome females exhibit lower fecundity (Banham 1990; Sánchez-Guillén et al. 2017) and more aggressive behaviors towards approaching males (Sánchez-Guillén et al. 2017). Although androchrome females show only a slightly lower number of copulations compared to the gynochrome females (Cordero 1987; 1992), male mimicry remains a key strategy for reducing mating frequency.

Our results extend this interpretation by showing that male mimicry in androchrome females of *I. elegans* also involves the morphology of the prothorax, a structure directly involved in tandem formation. Moreover, that morphological distinction between female morphs is more pronounced in certain contexts, with clear differences in allopatry and in the NC hybrid zone, but reduced in the NW hybrid zone where hybridization and introgression are more extensive (Arce-Valdés et al. 2025a). This spatial pattern suggests that hybrid zone dynamics such as introgression and gene flow may modulate the degree of morphological differentiation between morphs, which could in turn influence patterns of female polymorphism.

In *I. graellsii*, both female morphs differ from males in prothorax shape but remain similar to each other, while in prothorax size, the two morphs differ from each other, with males showing intermediate values. Importantly, these morphological patterns are consistent across allopatry and both hybrid zones, suggesting that female morph differentiation in this species is stable across geographic contexts. This slight and stable differentiation may indicate that female morphs experience more uniform selective pressures, with less pronounced divergence linked to mimicry or size-related strategies. This could indicate that, in *I. graellsii*, the role of male mimicry is less central, and sexual conflict over mating rates may be weaker. Moreover, the consistent morphology between female morphs across zones and the relatively stable color-morph frequencies observed in *I. graellsii* (e.g., Andrés et al., 2000; Fincke et al., 2005; Sánchez-Guillén et al. 2011b) suggest that female polymorphism in this species is maintained by relatively stable selective pressures, likely involving sexual selection through male mating preferences and balancing selection associated with sexual conflict over mating rates.

### Morph-specific reinforcement and asymmetric reproductive character displacement

Building on the morphological similarity between androchrome females and males described above, our study confirmed asymmetric RCD between species in the prothorax shape, as this pattern was detected only in *I. elegans* females from both hybrid zones and was absent in *I. graellsii*. Importantly, this pattern was detected in both female colour morphs, indicating that divergence in this reproductive trait affects both androchrome and gynochrome females. This result is consistent with the previous morphometric analyses of Ballén-Guapacha et al. (2024) who also detected RCD in prothorax shape in *I. elegans* females from both hybrid zones. In contrast, RCD in the size of the prothorax showed a more restricted pattern, as it was detected only in the gynochrome females of *I. elegans* from the NC hybrid zone.

First, our results confirm that differences in the detection of RCD between hybrid zones cannot be explained by differences in the proportion of androchrome and gynochrome females in our dataset, because we analysed each female morph separately. Specifically, RCD in prothorax shape was detected in both female morphs, whereas RCD in CS was detected only in gynochrome females. Instead, the absence of size related RCD in the NW hybrid zone is more consistent with the high levels of interspecific gene flow reported for this region (Arce-Valdés et al. 2025a), which can reduce morphological divergence between species (Ballén-Guapacha et al., 2024). Second, the asymmetric RCD in prothorax size between female morphs closely matches the pattern of asymmetric reinforcement between morphs in both species detected for the mechanical barrier involved in the tandem formation, which depends on the interaction between male caudal appendages and the female prothorax (Ordaz-Morales et al., 2025). In that study, gynochrome females experienced stronger selection against heterospecific mating attempts than androchrome females, producing morph specific reinforcement of mechanical isolation. Together, these results are consistent with the idea that reinforcement and RCD operate differently between female morphs in *I. elegans*.

This pattern can be explained by differences in the susceptibility of each female morph to heterospecific mating attempts. Experimental studies have shown that androchrome females from allopatric populations already exhibit strong mechanical isolation from heterospecific *I. graellsii* males, because their male like prothorax morphology reduces the mechanical fit required for tandem formation (Ordaz-Morales et al., 2025). This asymmetry is likely the result of gynochrome females experiencing a higher number of both intraspecific and interspecific copulation attempts compared to androchrome females (Cordero-Rivera & Andrés-Abad, 2001; Cordero-Rivera & Sánchez-Guillén, 2007; Sánchez-Guillén et al., 2017). Under this scenario, androchrome females may initially experience weaker selection from reinforcement because their male mimicking morphology already reduces the probability of successful heterospecific tandem formation (Ordaz-Morales et al., 2025). This initial advantage in sympatry of the androchrome females may help explain the slightly higher frequency of androchrome females in both hybrid zones compared to the surrounding allopatric regions for both species (see Sánchez-Guillén et al., 2005). In other polymorphic taxa, species exhibiting RCD can transition from producing both morphs to producing only the less similar morph (between species) or show different distributions in morph frequency (Rice & Pfennig, 2006). These findings support the hypothesis that androchrome male mimicry may contribute to reproductive isolation and reinforce prezygotic barriers, as proposed by Johnson (1975).

### Correlated divergence of male traits involved in the tandem formation

Our analyses extend previous evidence of RCD in male traits in these hybrid zones. We detected RCD in prothorax shape in *I. elegans* males from the NC hybrid zone. This pattern is consistent with previous evidence showing RCD in male caudal appendages in both species and hybrid zones (Ballén-Guapacha et al., 2024), suggesting that several components of the mechanical coupling system involved in tandem formation may evolve under reinforcement.

Although the male prothorax does not directly participate in clasping during tandem formation, male caudal appendages interact mechanically with the female prothorax (Robertson & Paterson, 1982; McPeek et al., 2008; 2009). The lock-and-key mechanism during tandem formation likely reinforces this matching, with male genital traits evolving to maintain compatibility with conspecific females while preventing successful heterospecific matings (see Ballén-Guapacha et al., 2024). Therefore, divergence in male clasping structures may indirectly promote correlated divergence in other male traits associated with the tandem position. Consistent with this idea, we detected a positive correlation between the size of male caudal appendages and the male prothorax, which may reflect correlated variation between these traits. While this pattern aligns with general scaling relationships (e.g., isometric or allometric growth) and does not necessarily imply coevolution it may still be relevant for reproductive isolation. Previous studies have shown that male caudal appendages mechanically interact with the female prothorax during tandem formation, and that their morphology correlates with female prothorax shape (McPeek et al., 2008; 2009). Our findings extend this evidence by demonstrating that male traits, the prothorax and caudal appendages, also covary, suggesting a limited pattern of morphological covariation that could influence mating compatibility.

This covariation in size takes on additional significance when considering female morphs. Androchrome females, which mimic males in both prothorax size and shape, exhibit stronger reproductive isolation in allopatry, as they are less likely to form heterospecific tandems with *I. graellsii* males in hybrid zones compared to gynochrome females (Ordaz-Morales et al. 2025). Thus, the similarity between male and androchrome female prothorax morphology, combined with the internal correlation between male traits, may contribute to differences in mechanical compatibility during tandem formation. These patterns are consistent with the lock-and-key mechanism escribed for these species, in which divergence in male and female structures maintains compatibility with conspecific partners while preventing successful heterospecific matings (see Ballén-Guapacha et al., 2024). Such pressures are crucial for preserving species boundaries in zones of secondary contact.

## Conclusion

In *I. elegans*, male-mimicry in androchromes is reflected in both size and shape of the prothorax, but hybridization reduces shape distinctions between morphs, while size-based RCD is asymmetric and concentrated in gynochromes. In contrast, *I. graellsii* shows weaker morphological differentiation and convergence between female morphs and males in the hybrid zones, suggesting species-specific dynamics in response to hybridization.

Our results indicate that female polymorphism may influence the evolutionary response to hybridization, promoting asymmetric divergence of reproductive traits between morphs and species, while reinforcement can generate morph-specific patterns of RCD and involve multiple components of the mechanical coupling system involved in tandem formation. These findings highlight how female polymorphism can shape the evolutionary dynamics of RCD and reinforcement in polymorphic damselflies.

## ACKNOWLEDGEMENTS

We thank “Grupo Zalandrana de Odonatología (ADEMAR-RIOJA)” for kindly helping with collecting and sending samples, and Adolfo Cordero Rivera who facilitated the use of space and material in his laboratory at the University of Vigo. We are grateful to Janet Nolasco Soto and Jesús Ramses Chávez Ríos for their technical support. AVB-G received a PhD grant from the Mexican CONACyT. The research was funded by the Mexican CONACYT grant to RAS-G (282922). Capture permits from the studied zones were granted to RAS-G.

## AUTHOR CONTRIBUTIONS

RAS-G designed the study. AVB-G and RAS-G did field samplings. Data were analyzed by AVB-G. AVB-G and RAS-G designed analyses. AVB-G and RAS-G wrote the first draft of the manuscript. AVB-G made scientific illustrations. All authors read, reviewed, and approved the final version of the submitted manuscript.

## REFERENCES

1. Adams D. C., Collyer M. L., Kaliontzopoulou A., Baken E. (2022). Geomorph: Software for geometric morphometric analyses. R package version 4.0.4. https://cran.r-project.org/package=geomorph.

2. Andrés, J. A., Sánchez-Guillén, R. A., & Rivera, A. C. (2000). Molecular evidence for selection on female color polymorphism in the damselfly *Ischnura graellsii*. Evolution, 54(6), 2156–2161.

3. Arce-Valdés, L. R., Swaegers, J., Ballén-Guapacha, A., Chávez-Ríos, J., Chauhan, P., Wellenreuther, M., Hansson, B. & Sánchez-Guillén, R. A., (2025a). Hybridization outcomes reflect context-dependent reproductive isolation in two damselfly hybrid zones. BioRxiv. doi: 10.64898/2025.12.23.696213

4. Arce-Valdés L.R., Ballén-Guapacha A.V., Rivas-Torres, A., Chávez-Ríos J.R., Wellenreuther M., Hansson B., Sánchez-Guillén R.A. (2025b). Testing the predictions of reinforcement: long-term empirical data from a damselfly mosaic hybrid zone. Journal of Evolutionary Biology, 38 (1), 10–27.

5. Ballen-Guapacha, A. V., Ospina-Garces, S. M., Guevara, R., & Sánchez-Guillén, R. A. (2024). Reproductive character displacement: insights from genital morphometrics in damselfly hybrid zones. Heredity, 133, 355–368.

6. Banham, W. M. T. (1990). Non-random mating in the polymorphic damselfly Ischnura elegans. PhD thesis. The University of Manchester (United Kingdom).

7. Barnard A. A., Fincke O. M., McPeek M. A., Masly J. P. (2017). Mechanical and tactile incompatibilities cause reproductive isolation between two young damselfly species. Evolution, 71(10), 2410–2427.

8. Barrett, R. D., & Schluter, D. (2008). Adaptation from standing genetic variation. Trends in Ecology & Evolution, 23(1), 38–44.

9. Brennan, P. L. (2016). Studying genital coevolution to understand intromittent organ morphology. Integrative and comparative biology, 56(4), 669–681.

10. Bookstein F. L. (1991). Thin-plate splines and the atlas problem for biomedical images. Biennial International Conference on Information Processing in Medical Imaging, 326–342.

11. Collyer M. L., Adams DC (2018). RRPP: An r package for fitting linear models to high-dimensional data using residual randomization. Methods in Ecology and Evolution 9(7), 1772–1779.

12. Collyer M. L., Adams DC (2021). RRPP: Linear model evaluation with randomized residuals in a permutation procedure. https://CRAN.R-project.org/package=RRPP.

13. Cooley, J. R. (2007). Decoding asymmetries in reproductive character displacement. Proceedings of the Academy of Natural Sciences of Philadelphia, 156(1), 89–96.

14. Cordero, A. (1987). Estructura de población en *Ischnura graellsii* Rambur, 1842 (Zygoptera: Coenagrionidae). Boletín de la Asociación española de Entomología, 11, 269–286.

15. Cordero, A. (1989). Reproductive behaviour of *Ischnura graellsii* (Rambur)(Zygoptera: Coenagrionidae). Odonatologica, 18(3), 237–244.

16. Cordero, A. (1990). The inheritance of female polymorphism in the damselfly *Ischnura graellsii* (Rambur) (Odonata: Coenagrionidae). Heredity, 64(3), 341–346.

17. Cordero, A. (1992). Density-dependent mating success and colour polymorphism in females of the damselfly *Ischnura graellsii* (Odonata: Coenagrionidae). Journal of Animal Ecology, 769–780.

18. Cordero-Rivera, A., & Andrés, J. A. (2001). Estimating female morph frequencies and male mate preferences of polychromatic damselflies: a cautionary note. Animal Behaviour, 61, F1–F6.

19. Cordero-Rivera, A., & Sánchez-Guillén, R. A. (2007). Male-like females of a damselfly are not preferred by males even if they are the majority morph. Animal Behaviour, 74(2), 247–252.

20. Cordero-Rivera, A., Rivas-Torres, A., & Sánchez-Guillén, R. A. (2024). Evolution in Islands: contrasting morph frequencies in damselfly populations of the Balearic Islands. Biological Journal of the Linnean Society, blad173.

21. Coyne, J. A., & Orr, H. A. (1989). Patterns of speciation in Drosophila. Evolution, 43(2), 362–381.

22. Dryden I. L., Mardia K. V. (2016). Statistical shape analysis with applications in R. John Wiley & Sons.

23. Fincke, O. M., Jödicke, R., Paulson, D. R., & Schultz, T. D. (2005). The evolution and frequency of female color morphs in Holarctic Odonata: why are male-like females typically the minority? International Journal of Odonatology, 8(2), 183–212.

24. Gosden, T. P., Stoks, R., & Svensson, E. I. (2011). Range limits, large-scale biogeographic variation, and localized evolutionary dynamics in a polymorphic damselfly. Biological Journal of the Linnean Society, 102(4), 775–785.

25. Gunz P., Mitteroecker, P. (2013). Semilandmarks: A method for quantifying curves and surfaces. Hystrix, 24, 103–109.

26. Hammers, M., & Van Gossum, H. (2008). Variation in female morph frequencies and mating frequencies: random, frequency-dependent harassment or male mimicry? Animal Behaviour, 76(4), 1403–1410.

27. Hammers, M., Sánchez-Guillén, R. A., & Van Gossum, H. (2009). Differences in mating propensity between immature female color morphs in the damselfly *Ischnura elegans* (Insecta: Odonata). Journal of insect behavior, 22, 324–337.

28. Hinnekint, B. O. N. (1987). Population dynamics of *Ischnura E. elegans* (Vander Linden) (Insecta: Odonata) with special reference to morphological colour changes, female polymorphism, multiannual cycles and their influence on behaviour. Hydrobiologia, 146, 3–31.

29. Hinojosa, J. C., Koubínová, D., Dincă, V., Hernández-Roldán, J., Munguira, M. L., García-Barros, E., … & Vila, R. (2020). Rapid colour shift by reproductive character displacement in Cupido butterflies. Molecular Ecology, 29(24), 4942–4955.

30. Hopkins, R. (2013). Reinforcement in plants. New Phytologist, 197(4), 1095–1103.

31. Howard D. J., & Harrison R. G. (1993). Reinforcement: origin, dynamics, and fate of an evolutionary hypothesis. In: Hybrid zones and the evolutionary process, (R. G. Harrison ed.), pp 46–69. Oxford University Press, New York.

32. Johnson, C. (1975). Polymorphism and natural selection in Ischnuran damselflies. Evol. Theory, 1, 81–90.

33. Joop, G., Siva-Jothy, M. T., & Rolff, J. (2006). Female colour polymorphism: gender and the eye of the beholder in damselflies. Evolutionary Ecology, 20, 259–270.

34. Kawano, K. (2002). Character displacement in giant rhinoceros beetles. The American Naturalist, 159(3), 255–271.

35. Klingenberg C. P. (2020). Walking on Kendall’s Shape Space: Understanding Shape Spaces and Their Coordinate Systems. Evol Biol 47: 334–352.

36. Liou L. W., Price TD (1994). Speciation by reinforcement of premating isolation. Evolution, 48, 1451–1459.

37. McPeek, M. A., Shen, L., & Farid, H. (2009). The correlated evolution of three-dimensional reproductive structures between male and female damselflies. Evolution, 63(1), 73–83.

38. McPeek, M. A., Shen, L., Torrey, J. Z., & Farid, H. (2008). The Tempo and Mode of Three-Dimensional Morphological Evolution in Male Reproductive Structures. The American Naturalist, 171(5), E158–E178.

39. McPeek M. A., Symes L. B., Zong D. M., McPeek C. L. (2011) Species recognition and patterns of population variation in the reproductive structures of a damselfly genus. Evolution, 65, 419–428.

40. Nava-Bolaños A., Sánchez-Guillén R. A., Munguía-Steyer R. (2017). Isolation barriers and genetic divergence in non-territorial Argia damselflies. Biological Journal of the Linnean Society 120, 804–817.

41. Norton, N. A., Fernando, M. T. R., Herlihy, C. R., & Busch, J. W. (2015). Reproductive character displacement shapes a spatially structured petal color polymorphism in Leavenworthia stylosa. Evolution, 69(5), 1191–1207.

42. Ordaz-Morales J. E., Juárez-Jiménez A. L., Stand-Pérez M., Arce-Valdés L. R., Ballen-Guapacha A., Chávez-Ríos J. R., Cordero-Rivera A., & Sánchez-Guillén R. A. (2025). Alternative reproductive strategies explain asymmetric reinforcement of reproductive isolation in two Ischnura damselfly species bioRxiv: https://www.biorxiv.org/cgi/content/short/2025.05.04.652146v1.

43. Pfennig, D. W., & Murphy, P. J. (2000). Character displacement in polyphenic tadpoles. Evolution, 54(5), 1738–1749.

44. Pfennig, D. W., & Murphy, P. J. (2002). How fluctuating competition and phenotypic plasticity mediate species divergence. Evolution, 56(6), 1217–1228.

45. Pfennig, K., & Pfennig, D. (2009). Character displacement: ecological and reproductive responses to a common evolutionary problem. The Quarterly review of biology, 84(3), 253–276.

46. R Core Team (2023). A language and environment for statistical computing. R Foundation for Statistical Computing, Vienna, Austria. URL https://www.R-project.org/.

47. Rice, A. M., & Pfennig, D. W. (2007). Character displacement: in situ evolution of novel phenotypes or sorting of pre-existing variation? Journal of Evolutionary Biology, 20(2), 448–459.

48. Robertson, H. M., & Paterson, H. E. (1982) Mate recognition and mechanical isolation in Enallagma damselflies (Odonata: Coenagrionidae). Evolution, 36(2), 243–250

49. Robertson, H. M. (1985). Female dimorphism and mating behaviour in a damselfly, *Ischnura ramburi*: females mimicking males. Animal Behaviour, 33(3), 805–809.

50. Rohlf F. J., Slice D. (1990). Extensions of the procrustes method for the optimal superimposition of landmarks. Systematic Zoology 39, 40–59.

51. Sánchez-Guillén, R. A., Van Gossum, H., & Cordero Rivera, A. (2005). Hybridization and the inheritance of female colour polymorphism in two ischnurid damselflies (Odonata: Coenagrionidae). Biological Journal of the Linnean Society, 85(4), 471–481.

52. Sánchez-Guillén, R. A., Wellenreuther, M., Cordero-Rivera, A., & Hansson, B. (2011a). Introgression and rapid species turnover in sympatric damselflies. BMC Evolutionary Biology, 11, 210.

53. Sánchez-Guillén, R. A., Hansson, B., Wellenreuther, M., Svensson, E. I., & Cordero-Rivera, A. (2011b). The influence of stochastic and selective forces in the population divergence of female colour polymorphism in damselflies of the genus *Ischnura*. Heredity, 107(6), 513–522.

54. Sanchez-Guillen, R. A., Wellenreuther, M., & Cordero Rivera, A. (2012). Strong asymmetry in the relative strengths of prezygotic and postzygotic barriers between two damselfly sister species. Evolution, 66(3), 690–707.

55. Sánchez-Guillén, R. A., Muñoz, J., Rodríguez-Tapia, G., Feria Arroyo, T. P., & Córdoba-Aguilar, A. (2013). Climate-induced range shifts and possible hybridisation consequences in insects. PloS one, 8(11), e80531.

56. Sánchez-Guillén R. A., Córdoba-Aguilar A., Cordero-Rivera A., Wellenreuther M. (2014). Rapid evolution of prezygotic barriers in non-territorial damselflies. Biological Journal of the Linnean Society 113(2), 485–496.

57. Sánchez-Guillén, R. A., Wellenreuther, M., Chávez-Ríos, J. R., Beatty, C. D., Rivas-Torres, A., Velasquez-Vélez, M., & Cordero-Rivera, A. (2017). Alternative reproductive strategies and the maintenance of female color polymorphism in damselflies. Ecology and Evolution, 7(15), 5592–5602.

58. Schlager S. (2017). “Morpho and Rvcg - Shape Analysis in R.” In Zheng G, Li S, Szekely G (eds.), Statistical Shape and Deformation Analysis, 217–256. Academic Press. ISBN 9780128104934.

59. Schluter, D. (2000). The ecology of adaptive radiation. OUP Oxford.

60. Sherratt, T. N. (2001). The evolution of female-limited polymorphisms in damselflies: a signal detection model. Ecology Letters, 4(1), 22–29.

61. Slatkin M. (1980). Ecological character displacement. Ecology, 61, 163–177.

62. Tripp, E. A., Dexter, K. G., & Stone, H. B. (2021). Reproductive character displacement and potential underlying drivers in a species-rich and florally diverse lineage of tropical angiosperms (Ruellia; Acanthaceae). Ecology and Evolution, 11(9), 4719–4730.

63. Wellenreuther M., Sánchez-Guillén R. A. (2016). Nonadaptive radiation in damselflies. Evolutionary Applications 9, 103–118.

64. Wellenreuther M., Muñoz J., Chávez-Ríos J. R., Hansson B., Cordero-Rivera A., Sánchez-Guillén R. A. (2018). Molecular and ecological signatures of an expanding hybrid zone. Ecology and Evolution 8, 4793–4806.

